# Expanding the genome information on *Bacillales* for biosynthetic gene cluster discovery

**DOI:** 10.1101/2024.04.24.590912

**Authors:** Lijie Song, Lasse Johan Dyrbye Nielsen, Xinming Xu, Omkar Satyavan Mohite, Matin Nuhamunada, Zhihui Xu, Rob Murphy, Kasun Bodawatta, Michael Poulsen, Mohamed Hatha Abdulla, Eva C. Sonnenschein, Tilmann Weber, Ákos T. Kovács

**Author notes:** Correspondence and requests for materials should be addressed to Ákos T. Kovács, and Tilmann Weber,. Contributed equally/Co-first authors (Lijie Song, Lasse Johan Dyrbye Nielsen).

## Abstract

This study showcases 121 new genomes of spore-forming *Bacillales* from strains collected globally from a variety of habitats, assembled using Oxford Nanopore long-read and MGI short-read sequences. *Bacilli* are renowned for their capacity to produce diverse secondary metabolites with use in agriculture, biotechnology, and medicine. These secondary metabolites are encoded within biosynthetic gene clusters (smBGCs). smBGCs have significant research interest due to their potential for the discovery of new bioactivate compounds. Our dataset includes 62 complete genomes, 2 at chromosome level, and 57 at contig level, covering a genomic size range from 3.50 Mb to 7.15 Mb. Phylotaxonomic analysis revealed that these genomes span 16 genera, with 69 of them belonging to *Bacillus*. A total of 1,176 predicted BGCs were identified by *in silico* genome mining. We anticipate that the open-access data presented here will expand the reported genomic information of spore-forming *Bacillales* and facilitate a deeper understanding of the genetic basis of *Bacillales*’ potential for secondary metabolite production.

## Background & Summary

*Bacillus* is a genus of Gram-positive, rod-shaped bacteria that are widely distributed in soil, water, and other diverse environments. *Bacillus* species have been extensively studied for their potential to produce secondary metabolites (SMs), which have a wide range of functions and activities, and are being harnessed in various fields, such as agriculture, biotechnology, and medicine^1,2^. Several studies have reported that *Bacillus* and related genera produce secondary metabolites, an ability conferred by the presence of biosynthetic gene clusters^3–5^.

Secondary metabolite biosynthetic gene clusters (smBGCs) are genomic regions containing two or more genes involved in the biosynthetic pathway of secondary metabolites. These genes encode enzymes, transport proteins, regulatory factors, and other accessory proteins that contribute to the secondary metabolite biosynthetic process^6^. The composition and structures of smBGCs can vary widely across and even within the same species. The importance and feasibility of exploring species-specific BGCs have been recently highlighted^7,8^. Many bioinformatics tools have been developed to predict, identify, and characterize smBGCs^9^, which require high quality genome sequences^10^. The development of sequencing technologies has made whole genome sequencing simpler and faster. In particular, the integration of high throughput sequencing (short-read) and long-read sequencing data, can lead to high quality assemblies of genomes, including complete genomes^11^.

In this study, we performed whole genome sequencing for strains collected from different countries and regions spanning four different continents (Online-only Figure 1), based on an integrated approach, including Oxford Nanopore long-read sequencing and MGI short-read sequencing. Here, we sequenced and assembled 121 genomes using this approach. An outline of the study’s experimental and analysis design is presented in Figure 1, and detailed descriptions of the workflow are provided in the methodology sections. According to the completeness criteria of the National Center for Biotechnology Information (NCBI), we produced, in total, 62 assemblies at a complete genome level, 2 at chromosome level, and a remaining 57 at contig level (Online-only Table 1 for details). Overall, the genome sizes range from 3.50 Mb to 7.15 Mb (5.09 Mb on average), with a GC content ranging from 34.50% to 54.00% (40.19% on average). Based on NCBI PGAP^12^, an average of 5,119 genes, including 4,851 protein-coding genes were annotated in the genomes (Table 1).

**Table 1.** Summary of general genome information for each genus

| *Genus* | # of isolates | Assembly level |  |  | Average genome size (min and max) (Mbps) | Average GC (min and max) (%) | Average gene number (min and max) |
| --- | --- | --- | --- | --- | --- | --- | --- |
|  |  | complete genome | chromosome | contig |  |  |  |
| *Bacillus* | 69 | 35 | 2 | 32 | 4.86<br>(3.50-6.33) | 39.46<br>(34.40-46.50) | 4,998<br>(3,616-6,554) |
| *Peribacillus* | 16 | 7 | 0 | 9 | 5.59<br>(5.35-6.02) | 40.25<br>(39.50-40.50) | 5,452<br>(5,245-5,817) |
| *Neobacillus* | 8 | 6 | 0 | 2 | 6.05<br>(5.59-6.29) | 38.63<br>(38.00-40.00) | 5,917<br>(5,383-6,233) |
| *Paenibacillus* | 5 | 3 | 0 | 2 | 6.40<br>(5.29-7.15) | 46.60<br>(44.00-50.00) | 5,724<br>(4,881-6,560) |
| *Lysinibacillus* | 5 | 2 | 0 | 3 | 5.17<br>(4.56-5.54) | 36.80<br>(36.50-37.50) | 5,180<br>(4,575-5,605) |
| *Cytobacillus* | 4 | 2 | 0 | 2 | 5.03<br>(4.73-5.30) | 40.50<br>(37.00-42.00) | 5,034<br>(4,752-5,251) |
| *Fictibacillus* | 4 | 1 | 0 | 3 | 5.90<br>(3.94-5.40) | 42.88<br>(39.50-44.00) | 5,070<br>(4,156-5,551) |
| *Brevibacillus* | 2 | 1 | 0 | 1 | 5.68 and 6.08 | 52.00 and 54.00 | 5,498 and 5,699 |
| *Mesobacillus* | 1 | 1 | 0 | 0 | 4.80 | 43.00 | 4,782 |
| *Siminovitchia* | 1 | 1 | 0 | 0 | 3.74 | 44.00 | 3,727 |
| *Halobacillus* | 1 | 1 | 0 | 0 | 3.68 | 47.00 | 3,841 |
| *Virgibacillus* | 1 | 1 | 0 | 0 | 4.01 | 37.50 | 3,880 |
| *Priestia* | 1 | 1 | 0 | 0 | 3.82 | 38.00 | 3,985 |
| *Rossellomorea* | 1 | 0 | 0 | 1 | 4.33 | 48.00 | 4,467 |
| *Ureibacillus* | 1 | 0 | 0 | 1 | 4.45 | 36.00 | 4,240 |
| *Ferdinandcohnia* | 1 | 0 | 0 | 1 | 4.77 | 37.50 | 4,746 |
| **In total** | **121** | **62** | **2** | **57** | **5.09**<br>**(3.50-7.15)** | **40.19**<br>**(34.50-54.00)** | **5,119 (3,616-6,560)** |

**Figure 1.**
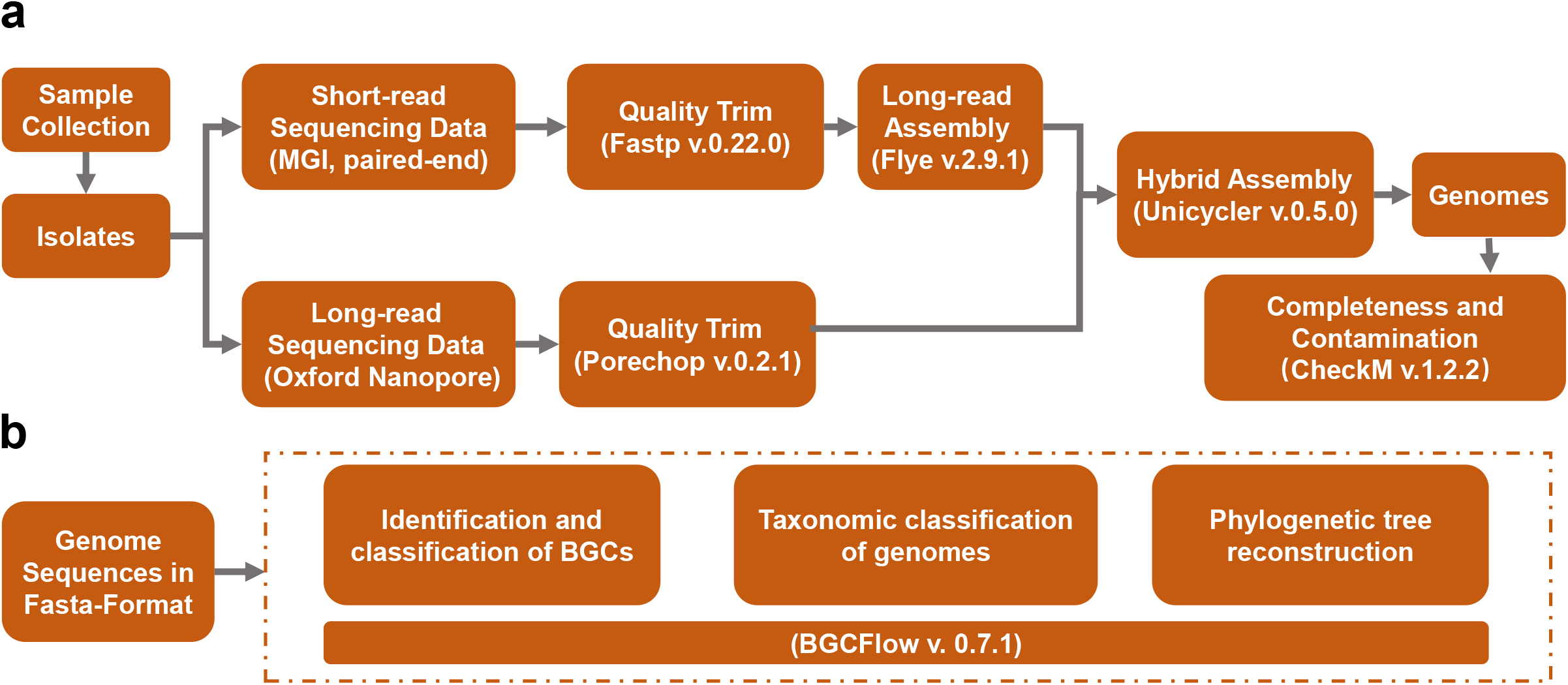
Illustration of Genome assembly and BGC analysis. (a) Strategy for sequencing and genome assembly, (b) the BGC analysis pipeline.

Taxonomic analysis showed that these 121 genomes could be classified into 16 genera within the *Bacillales* order, most of which were species from the *Bacillus* genus (Figure 2). (Online-only Table 2).

**Figure 2.**
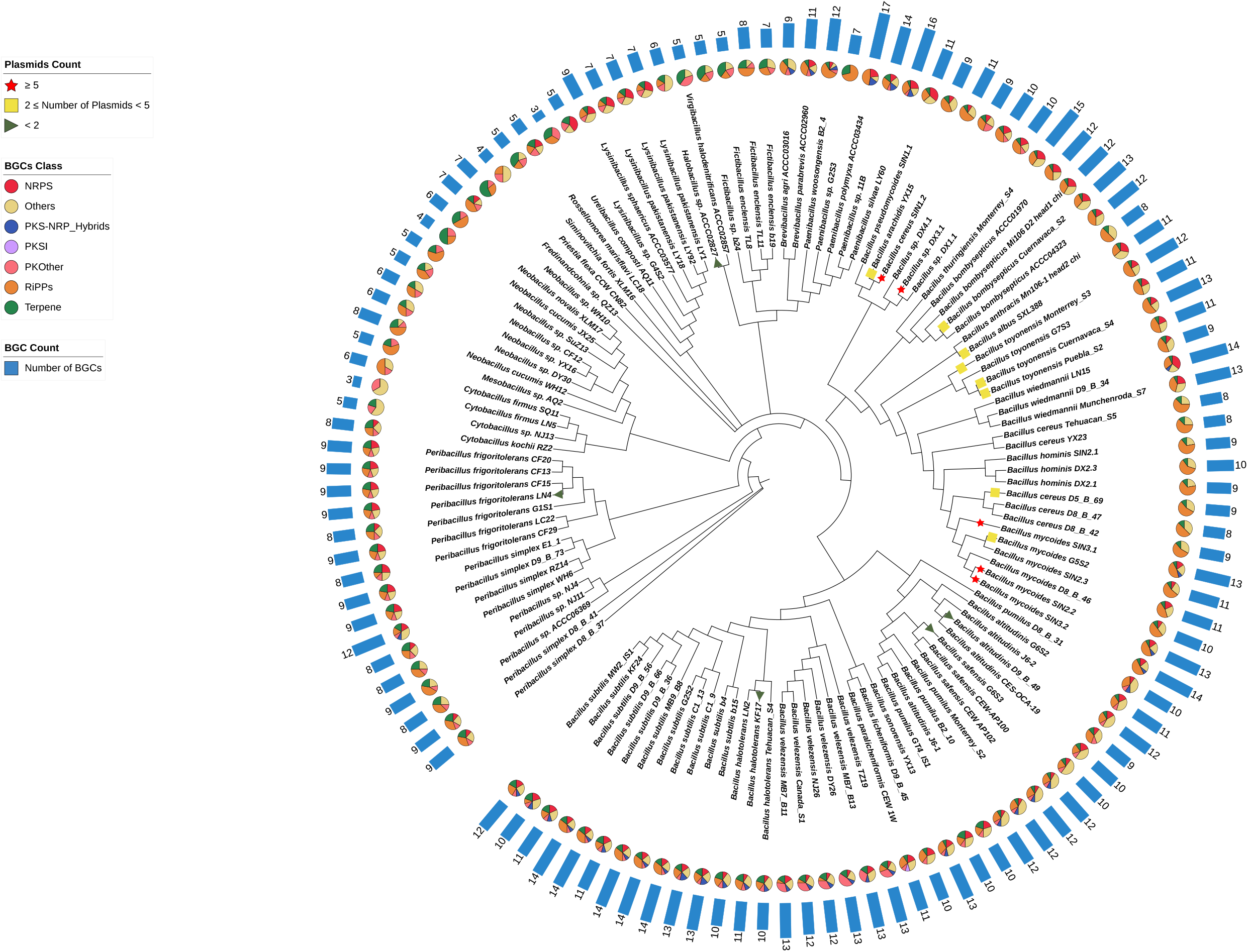
The phylogenetic trees of 121 genomes with plasmid content and BGC class and count indicated.

To assess the potential for secondary metabolite production in these isolates, the genome mining tool BGCFlow^13^ was applied for BGC identification and annotation, resulting in a total of 1,176 BGCs predicted. The BGCs were categorized into seven classes through BiG-SCAPE^14^, part of the BGCFlow executable, which showed that RiPPs have the greatest count of 381 and comprise the highest percentage at 32.4% (Online-only Table 3). The distribution of BGC counts per genus highlights the uneven abundance of BGCs between the distinct genera (Figure 3). Notably, the genera *Bacillus* and *Paenibacillus* harbor the highest number of BGCs among the genomes presented here.

**Figure 3.**
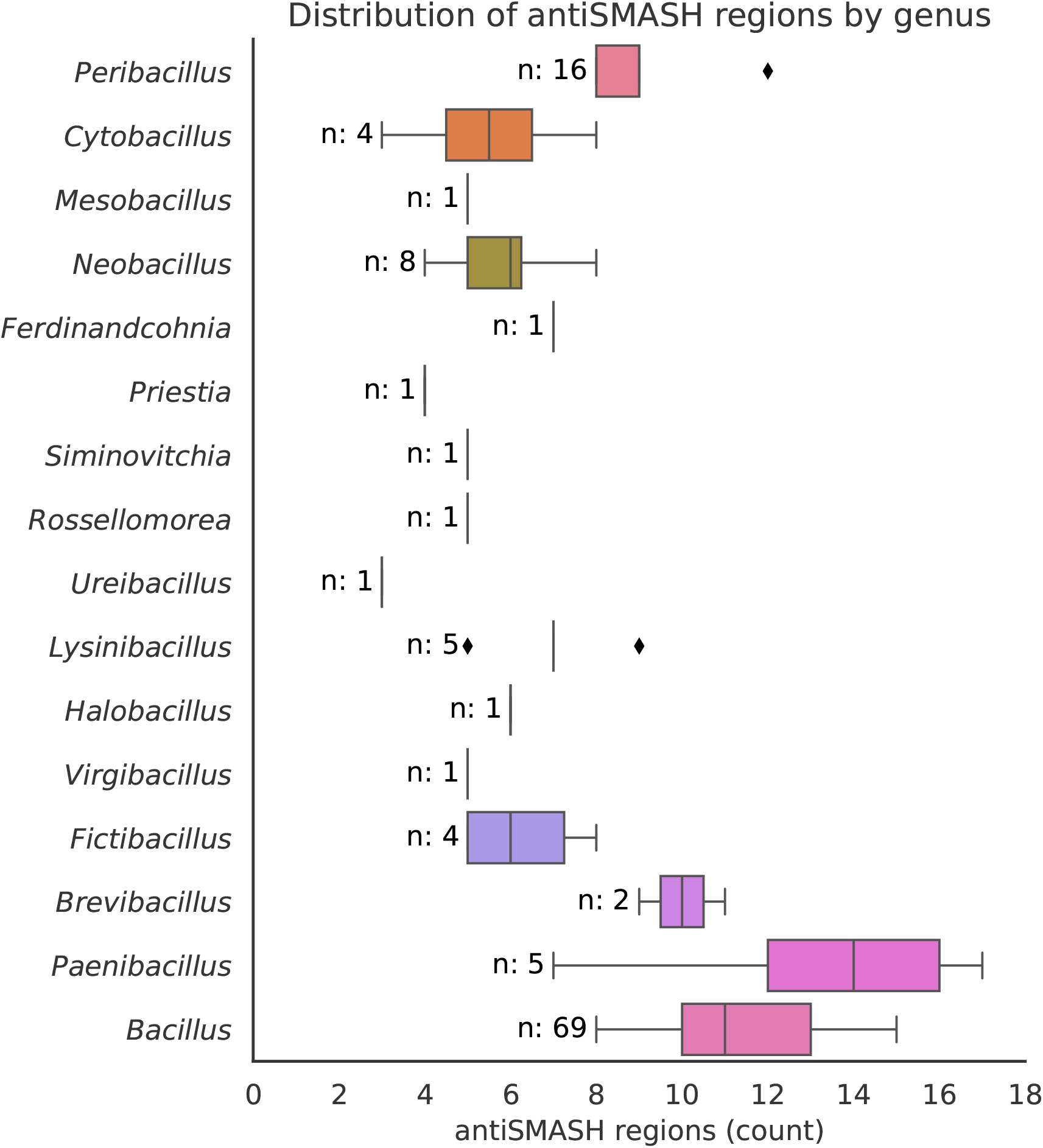
The number of BGCs in each genus of the 121 genomes.

The datasets and genomic analysis results described here greatly expand the reported genomic information of spore-forming *Bacillales* and will also strengthen studies advancing our understanding of the secondary metabolite potential of the *Bacillales* order.

## Method

### Sample collection and isolation

Sample collection was dependent on the isolating laboratory. Using soil samples collected at diverse locations in Germany, Denmark, China, and Mexico, spore-forming soil bacteria were isolated after heat treatment at 80°C for 10 minutes and spreading the soil suspension on lysogeny broth (LB) or tryptic soy broth (TSB) plates with 1.5% agar that were incubated at 37°C for 2 days.

*Bacillus altitudinis* J6-1 and J6-2 were isolated from a biofilm sample obtained from the pier at Jyllinge Harbour (55.744923; 12.094888). Biofilm samples were incubated at 80°C for 15 mins and subsequently plated on LB agar and incubated at 25°C.

Other marine samples were collected from the Cochin estuary and adjacent coastal waters (South-west coast of India), during pre-monsoon (March), monsoon (August) and post-monsoon (December) periods of the year 2012 and 2013. Water samples were serially diluted and spread on Norris Glucose Nitrogen free medium (NGNF medium, HIMEDIA-M712) with 1.5% agar (Himedia GRM 666) and incubated at 28±1 °C for 7-14 days. Separated colonies with different morphologies were picked using a sterile inoculation loop, restreaked and maintained on the slants of fresh nitrogen free culture medium at 4 °C. Cell morphology and presence of endospore was analyzed by light microscopy (Olympus CX21i). Rod shaped endospore forming isolates were selected for this study.

Isolate Mi106 D2 head1 chi was obtained from the head of a worker termite from a colony of *Microtermes* sp. and Mn106-1 head2 chi was obtained from the head of a worker from a colony of *Macrotermes natalensis* in Mookgophong, South Africa (S24 40 30.5 E28 47 50.4) in 2010. In both cases, the surface of a worker termite was rinsed using phosphate buffer saline (PBS). Subsequently, the head of the termite was crushed in 200 µl PBS, which was subsequently spread onto chitin medium (4 g chitin, 0.7 g K2HPO4, 0.3 g KH2PO4, 0.5 g MgSO4 × 5H2O, 0.01 g FeSO4 × 7H20, 0.001 g ZnSO4, 0.001 g MnCl2, and 20 g of agar per liter). Growing colonies on plates were streaked onto Yeast Malt Extract Agar medium (4 g yeast extract, 10 g malt extract, 4 g D-glucose and 20 g bacteriological agar per liter), and once in pure culture, stored in 10 % glycerol at −20 °C. Isolate 11B was obtained using the same approach on a fragment from a fungus garden of a *Macrotermes natalensis* colony collected in Rietondale, South Africa (S25 43 45.6 E28 14 09.9) in 2010.

Strains GT4_IS1 and MW2_IS1 were previously isolated from the uropygial glands of Great tits (*Parus major*) from Denmark and Czechia respectively^15^.

In each case, observed colonies were re-streaked to obtain single colonies, and subsequently stored at - 80°C with 28% glycerol added. To obtain primary information about these strains, colony PCR was employed to amplify the 16S rRNA gene. Strains that exhibited low similarity and distant branches in the 16S rRNA phylogenetic tree were selected for further study.

### Genomic DNA (gDNA) extraction

For genomic DNA (gDNA) extraction, a pure single colony of each isolate was inoculated in 5 ml of LB and incubated at 37°C for more than 12 hours. Then gDNA was extracted using E.Z.N.A. DNA extraction kits (OMEGA Bio-Tek Inc., Norcross, GA, USA) following the manufacturer’s instructions. The quality and quantity of gDNA were assessed using agarose gel electrophoresis and Nanodrop (Thermo Fisher Scientific, MA, USA), to guarantee that the integrity, concentration, and purity met the requirements for library construction and sequencing.

### Short-read sequencing on MGI platform

For each strain, 300 ng gDNA was used for short-read sequencing library construction according to MGI paired-end libraries construction protocol^16^. Briefly, gDNA was fragmented to 200-300 bp using segmentase followed by fragment selection with VAHTS™ DNA Clean Beads (Vazyme, Nanjing, Jiangsu, China). Subsequently, end repair, A-tailing reactions and adapter ligation were implemented. After PCR and purification, the concentration of each library was determined using Qubit® dsDNA HS Assay Kit (Thermo Fisher Scientific) as quality control. The qualified libraries were sequenced on the DNBSEQ-G400 (MGI Tech Co., Ltd.) platform according to the manufacturer’s instructions to generate paired end reads (150 bps at each end).

### Long-read sequencing on Oxford Nanopore platform

For Oxford Nanopore sequencing, the libraries were prepared using the SQK-RBK110.96 barcoding kit (Oxford Nanopore Technologies, Oxford, UK) starting from 50ng DNA for each strain. In brief, each sample was fragmented and ligated by a unique rapid barcode with incubation at 30°C for 2 minutes and then at 80 °C for 2 minutes, then all barcoded samples were pooled together in a 1:1 ratio and purified by SPRI beads. After ligation of 1 µl of Rapid Adapter F (RAP F) to 11 µl of pooled DNA, the final library was quantified using Nanodrop. The ONT library was loaded into the MinION spot-on Flow Cell (R9 Version) and sequenced on a MinION Mk1B device according to standard protocol. The resulting reads were base called and demultiplexed with MinKNOW UI v.4.1.22.

### Genome assembly

For *de novo* assembly, the MGISEQ paired end short reads were adapter and quality trimmed using fastp v.0.22.0 and the Nanopore long reads were adapter trimmed using porechop v.0.2.1, using standard settings^17,18^. The trimmed long reads from Nanopore were assembled with flye v.2.9.1-b1780, and subsequently the trimmed reads from both platforms and the long-read assembly were hybrid assembled with Unicycler v.0.5.0 using the *–existing_long_read_assembly* option^19,20^. The completeness and contamination levels of each strain were assessed using CheckM v.1.2.2^21^.

### Genome annotation, taxonomic analysis and BGC prediction

The genomes of the 121 isolates were taxonomically classified and gene-annotated in a two-step process. Initially, we employed GTDB-Tk v2.11, using the ‘classify_wf’ command, to preliminarily assign taxonomic classifications to the FASTA format genomes. Subsequently, these genomes were uploaded to the NCBI GenBank database, where they were annotated using the NCBI Prokaryotic Genome Annotation Pipeline (PGAP)^12,22^. Following this, we conducted a comprehensive analysis of the annotated genomes using BGCFlow v0.7.1. This tool integrates multiple genome mining and phylogenetic tools into one pipeline^13^. To set up the analysis, we created a folder containing the project configuration structure as defined by BGCFlow Portable Encapsulated Project (PEP)^23^ specification. The designated project folder contains a comma separated sample file which contains the NCBI-assigned GenBank accession numbers of the 121 de novo assembled genomes and the PEP configuration file for the BGCFlow run. The YAML configuration file for the project was configured to enable GTDB-Tk and autoMLST wrapper for phylogenetic tree construction, antiSMASH^24^ for BGC annotation, and BiG-SCAPE^14^ for BGC dereplication and generating summary tables. BGCFlow was executed using standard settings.

We conducted a non-exhaustive search for plasmids within our *de novo* assembled genomes by identifying contiguous sequences (contigs) as plasmids if they were circular and if RFPlasmid^25^ (v.0.0.18), an open-source software that classifies contigs as plasmid or chromosomal based on the presence of marker genes and k-mers, classified them as plasmids. Due to the incomplete assembly of several genomes, which resulted in the presence of linear fragments, the absence of any plasmid identified by this method does not necessarily indicate their true absence.

## Supporting information

Oline only Table 1

Oline only Table 2

Oline only Table 3

Oline only Table 4

Oline only Table 5

## Data Records

The sample information, assembled genomes, and raw reads of long-read sequencing on Nanopore and short-read sequencing on MGISEQ have been deposited in NCBI at BioProject under PRJNA960711 (https://www.ncbi.nlm.nih.gov/bioproject/PRJNA960711) (Online-only Table 1 and Online-only Table 4 for accession and other details).

## Technical Validation

In this study, the main steps of experimental procedures and data analysis have been validated. For short-read sequencing on MGI, the libraries were quantified with a minimum of 10 ng/μl. For *de novo* assembly, default parameters were used for quality trimming. In brief, after filtering, an average of 2.69 G MGI reads (0.66 G-6.52 G, PE150) and 76,507 Nanopore reads with mean N50 of 6,709 bp (1,777bp-13,698bp) for each sample were generated (Online-only Table 5). CheckM was used for validation of the genome completeness and contamination.

## Usage Notes

Not used

## Code availability

The software versions and parameters used for sequence filtering, assembly, and genome mining in this work are described in Methods. Custom code for setting up the BGCFlow run, processing the output, and producing figures, as well as for downloading the genomes, is available at https://github.com/ljdnielsen/bacillales_genomes_figures; https://doi.org/10.5281/zenodo.10907189.

## Acknowledgements

This project was supported by the Danish National Research Foundation (DNRF137) for the Center for Microbial Secondary Metabolites, and Novo Nordisk Foundation within the INTERACT project of the Collaborative Crop Resiliency Program (NNF19SA0059360). TW acknowledges funding from the Novo Nordisk Foundation Center for Biosustainability (NNF20CC0035580)

## Author Contributions

LS performed MGISEQ and Nanopore sequencing, analysis of genomes, interpreted the data, and wrote the manuscript.

LJDN performed Nanopore sequencing, assembly, and analysis of genomes, interpreted the data, and wrote the manuscript.

XX provided bacterial isolates, performed 16S rRNA gene sequencing, preliminary 16S rRNA-based phylotaxonomics, data visualization and helped to write the manuscript.

OSM helped with data analysis, contributed with BGCFlow, and helped to write the manuscript.

MN contributed with BGCFlow and helped to write the manuscript.

ZX provided new bacterial isolates and helped to write the manuscript.

RM provided new bacterial isolates and helped to write the manuscript.

KB provided new bacterial isolates and helped to write the manuscript.

MP provided new bacterial isolates and helped to write the manuscript.

MHA provided new bacterial isolates and helped to write the manuscript.

ECS provided new bacterial isolates and helped to write the manuscript.

TW conceived and supervised the project, contributed with BGCFlow, and wrote the manuscript.

ÁTK, conceived and supervised the project, and wrote the manuscript.

## Figures legends

**Online-only Figure 1** Distribution of sample collection site coordinates depicted using OpenStreetMap.

## Tables legends

**Online-only Table 1**. Summary characteristics of genome assembly and annotation

**Online-only Table 2**. Taxonomic placement of the 121 genomes

**Online-only Table 3**. Summary statistics of the BGCs in the 121 genomes

**Online-only Table 4**. Datasets on the 121 isolates

**Online-only Table 5**. Sequencing data quality of the 121 isolates

---

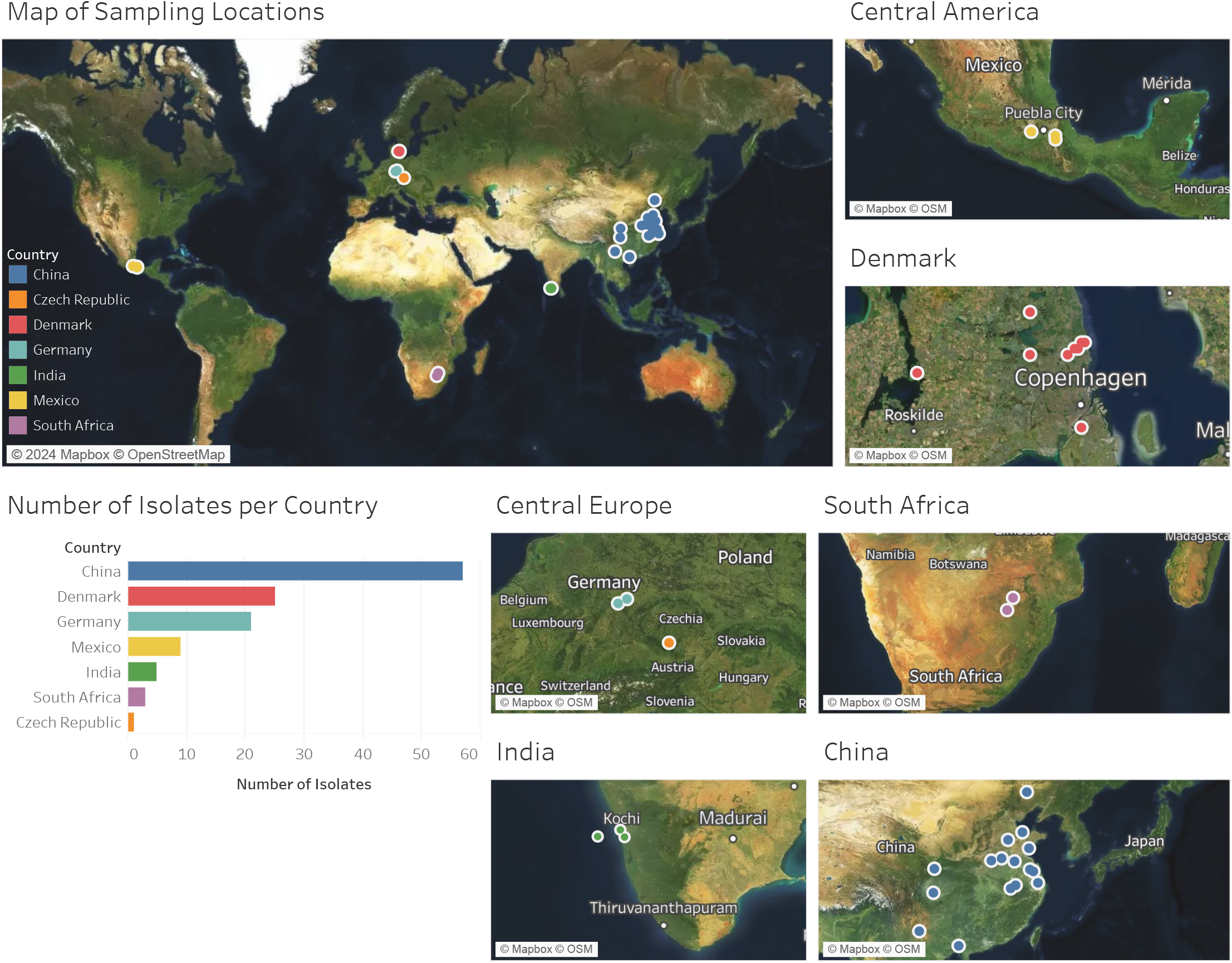

